# Chiral Histidine-Modified Gold Nanoclusters Loaded into Cationic Lipid Nanoparticles for Treatment of Biofilm- associated Infections

**DOI:** 10.64898/2026.02.25.707929

**Authors:** Zijian Ye, Xin Jin, Arnold Koekman, Mies van Steenbergen, Yang Liu, Zhentao Xing, Cornelis Seinen, Azin Khodaei, Enrico Mastrobattista, Joost PG Sluijter, Harrie Weinans, Raymond M. Schiffelers, Jaqueline Lourdes Rios, Bart van der Wal, Zhiyong Lei

**Affiliations:** Department of Orthopedics, University Medical Center Utrecht, Utrecht 3584 CX, The Netherlands; Laboratory of CDL Research, University Medical Center Utrecht, Utrecht 3508 GA, The Netherlands; Department of Pharmaceutics, Utrecht Institute for Pharmaceutical Sciences, Utrecht University, Utrecht 3584 CG, The Netherlands; College of Chemistry and Chemical Engineering, Central South University, Changsha, Hunan 410083, China; Department of Urology, University Medical Center Utrecht, Utrecht 3584 CX, The Netherlands; Laboratory of Experimental Cardiology, Department Heart & Lungs, University Medical Center Utrecht, Utrecht 3584 CX, The Netherlands; UMC Utrecht Regenerative Medicine Center, Circulatory Health Research Center, University Medical Center Utrecht, Utrecht University, Utrecht 3508 GA, The Netherlands; Department of Biomechanical Engineering, Faculty of Mechanical Engineering, Delft University of Technology, Delft 2628 CD, The Netherlands; Department of Orthopedics, Leiden University Medical Center, Leiden 2333 ZA, The Netherlands

**Author notes:** Corresponding authors: Zhiyong Lei; Bart C. H. van der Wal; Jaqueline Lourdes Rios. These authors contributed equally and share first authorship (Zijian Ye, Xin Jin).

**Keywords:** Antimicrobial resistance, Lipid nanoparticle, Nanocluster, Nanomedicine, Biofilm, Infection, Staphylococcus aureus

## Abstract

Antimicrobial resistance has heightened the risk of device-associated infections, in which *Staphylococcus aureus* (*S. aureus*) biofilms persist through extracellular-matrix protection and marked physiological heterogeneity. Gold nanoclusters (AuNCs) can disrupt biofilms via redox-associated stress, but limited in vivo stability and poor local retention constrain their translational potential. Here, we synthesized chiral histidine–modified AuNCs (D-, L-, and DL-AuNCs) and formulated them into mildly cationic lipid nanoparticles (AuNCs@LNP) by microfluidic assembly to enhance biofilm engagement while preserving nanocluster identity. The resulting particles were ∼100–120 nm with high gold loading and a modest charge reversal. In established *S. aureus* USA300 biofilms, free DL-AuNCs reduced viable burden by up to 5.6 log10 CFU/mL, suppressed metabolic activity, and increased ROS signaling. Notably, LNP encapsulation maintained bactericidal activity and further improved killing by ∼1 log10 CFU at matched Au dose, while substantially enhancing biomass removal (crystal violet residual biomass: 63.77% vs 45.17%), consistent with carrier-mediated matrix destabilization. In a subcutaneous implant infection model, a single perilesional dose of DL-AuNCs@LNP reduced implant-associated bacterial burden by 2.6 log10 CFU per implant versus PBS. These results establish a modular antibiofilm platform that couples nanocluster-driven killing with lipid-facilitated biofilm disruption and defines an efficacy–tolerability window to guide optimization of locally delivered antibiofilm nanomedicines.

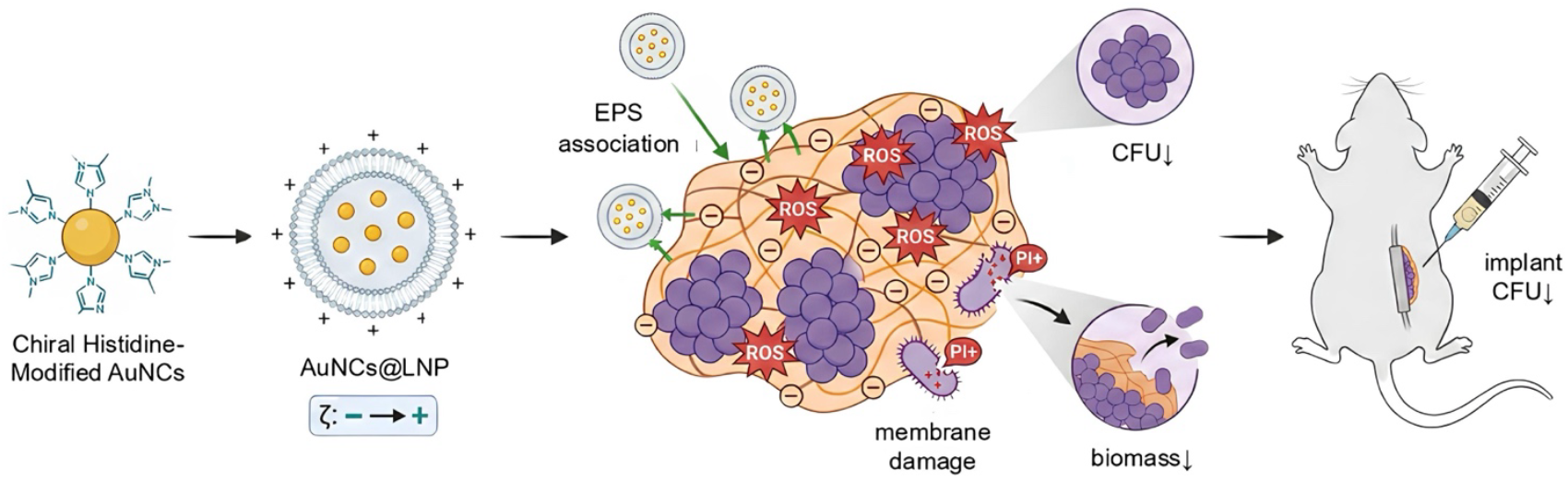

## 1. Introduction

Antimicrobial resistance (AMR) is steadily weakening modern medicine by allowing infections that are usually manageable to persist, recur, and disseminate. In the absence of effective countermeasures, projections point to an increasing global mortality and healthcare burden (1,2). Among AMR pathogens, Staphylococcus aureus (S. aureus), especially methicillin-resistant *S. aureus* (MRSA), continues to cause severe disease in both healthcare and community settings (3,4). It is also a prominent contributor to device- and implant-associated infections, which are notoriously difficult to eradicate (3,4). Importantly, the challenge is not limited to resistance against particular antimicrobial classes; *S. aureus* can shift into protective growth states that lower antimicrobial susceptibility and enable escape from host immune clearance.

The distinguishing feature of recalcitrant *S. aureus* infections is biofilm formation. Biofilms are essentially structured communities of microorganisms that are encapsulated in a self-secreted extracellular polymer matrix (EPS) capable of adhering to tissue surfaces and surfaces of biomaterials (5). Spatially heterogeneous nutrient and oxygen gradients exist within biofilms, and bacteria often shift to a hypometabolic or quasi-dormant phenotype as a result, a state that significantly reduces the efficacy of growth-dependent antibiotics (6). EPS matrices further impede the penetration of antimicrobial drugs, which can trap or neutralize drug molecules and form a protective microenvironment, and as a result, both the drug concentration and the exposure time needed to achieve bactericidal efficacy compared to planktonic state are significantly increased (7–9). In addition, biofilms enrich subpopulations of resident bacteria and promote horizontal gene transfer, accelerating the evolution of tolerance and drug resistance (10,11). These properties explain the failure of standard therapeutic regimens to clear biofilm reservoirs, ultimately leading to recurrence of infection and repeated clinical interventions.

Nanomaterials present promising approaches for countering biofilm-mediated protection by leveraging physicochemical characteristics. Metal nanoclusters (NCs) exhibit high surface-area-to-volume ratios alongside tunable surface chemistries, which enables diverse antimicrobial mechanisms to operate in concert (12–15). A substantial number of NC platforms are capable of generating reactive oxygen species (ROS) and triggering oxidative stress, which subsequently damages membranes, proteins, and nucleic acids—this mode of action yields broad-spectrum antibacterial effects with reduced reliance on any single biochemical target (16–21). Gold nanoclusters (AuNCs) have shown particular attraction for clinical translation due to their superior biocompatibility characteristics relative to other metallic nanomaterials, as well as their easily functionalized and modified surfaces, which modulate stability and biological interactions (22,23). In parallel, nanoscale chirality has emerged as a design parameter: chiral ligands can influence nano–bio interactions, including engagement with bacterial envelopes, and may alter antimicrobial performance in context-dependent manners (24,25).

Despite these compelling antimicrobial properties of gold nanoclusters, they face several practical challenges. Under physiological conditions, AuNCs may suffer from insufficient colloidal stability, which reduces their accessible surface area and weakens their ROS-dependent antimicrobial effect (26). In addition to maintaining their intrinsic activity, successful removal of formed biofilms requires not only bacterial killing, but also adequate retention and uniform distribution in the biofilm microenvironment, and these requirements are often compromised by rapid dispersion or removal after local delivery (27–29). Therefore, delivery strategies must be developed to maintain the antimicrobial capacity of AuNCs while enhancing their binding to biofilm structures and prolonging their functional presence at the biofilm-tissue interface.

Lipid nanoparticles (LNPs) are a clinically validated and compositionally diverse carrier platform that can address many of the limitations mentioned above (30–33). LNPs protect the encapsulated payload by tuning lipid composition and surface properties. They can also improve colloidal stability and modulate bioadhesion and tissue binding through charge adjustment, ultimately increase the bioavailability. This is particularly important in the field of biofilm therapeutics because the major extracellular polymer components typically carry a net negative charge. Thus, cationically or positively charged nanoparticle surfaces may facilitate electrostatic binding to the matrix, thereby promoting localized aggregation and disrupting matrix integrity (30–33). Although LNP has been widely developed for various biomedical applications, a key mechanistic and translational question remains for anti-biofilm therapies: can LNP vectors act not only as passive delivery carriers but also as active biofilm-binding components that synergize with the antimicrobial core of gold nanoclusters while avoiding excessive localized inflammatory responses (34,35)? Addressing this question requires a systematic side-by-side evaluation of free AuNCs with LNP-encapsulated AuNCs through comprehensive physicochemical analyses, complementary anti-biofilm assays, and in vivo validation using biofilm-associated infection models.

Here, we develop a modular nanomedicine that couples chiral histidine–modified gold nanoclusters with a cationic lipid nanoparticle platform (AuNCs@LNP) to improve treatment of *S. aureus* biofilm–associated infections. Histidine was chosen as a compact, biologically relevant ligand that stabilizes AuNCs while allowing direct comparison of chiral variants (D-, L-, and DL-). We hypothesize that AuNCs act as a potent antibacterial core via oxidative stress, and that encapsulation within cationic LNPs increases interactions with the anionic biofilm matrix, thereby improving functional engagement with biofilm architecture while maintaining AuNCs activity. To test these hypotheses, we (i) establish comprehensive physicochemical profiles of free AuNCs and AuNCs@LNP, including size, dispersity, surface charge, optical properties, and encapsulation efficiency; (ii) quantify antibiofilm activity against *S. aureus* USA300 using complementary readouts—biomass, viable burden, metabolic activity, oxidative stress, and structural integrity— to distinguish bactericidal effects from biomass disruption; and (iii) evaluate therapeutic performance and tolerability in a murine subcutaneous implant biofilm model after local administration, defining dose- and formulation-dependent constraints that are likely to be critical for clinical translation.

## 2. Results and Discussion

### 2.1 Cationic biofilm-interaction interface with preserved nanocluster identity

We established a formulation workflow that preserves AuNCs characteristics while enabling systematic tuning of the nano–bio interface (Figure 1A). Chiral histidine-stabilized Au nanoclusters were synthesized using histidine enantiomers as compact, biologically relevant ligands that also serve as mild reductants, allowing direct comparison of D-, L-, and DL-variants. Transmission electron microscopy (TEM) revealed dispersed ultrasmall monodispersed electron-dense features (Figure 1B, top) and did not show signatures indicative of growth into plasmonic AuNCs within the resolution and sampling of the analysis.

**Figure 1.**
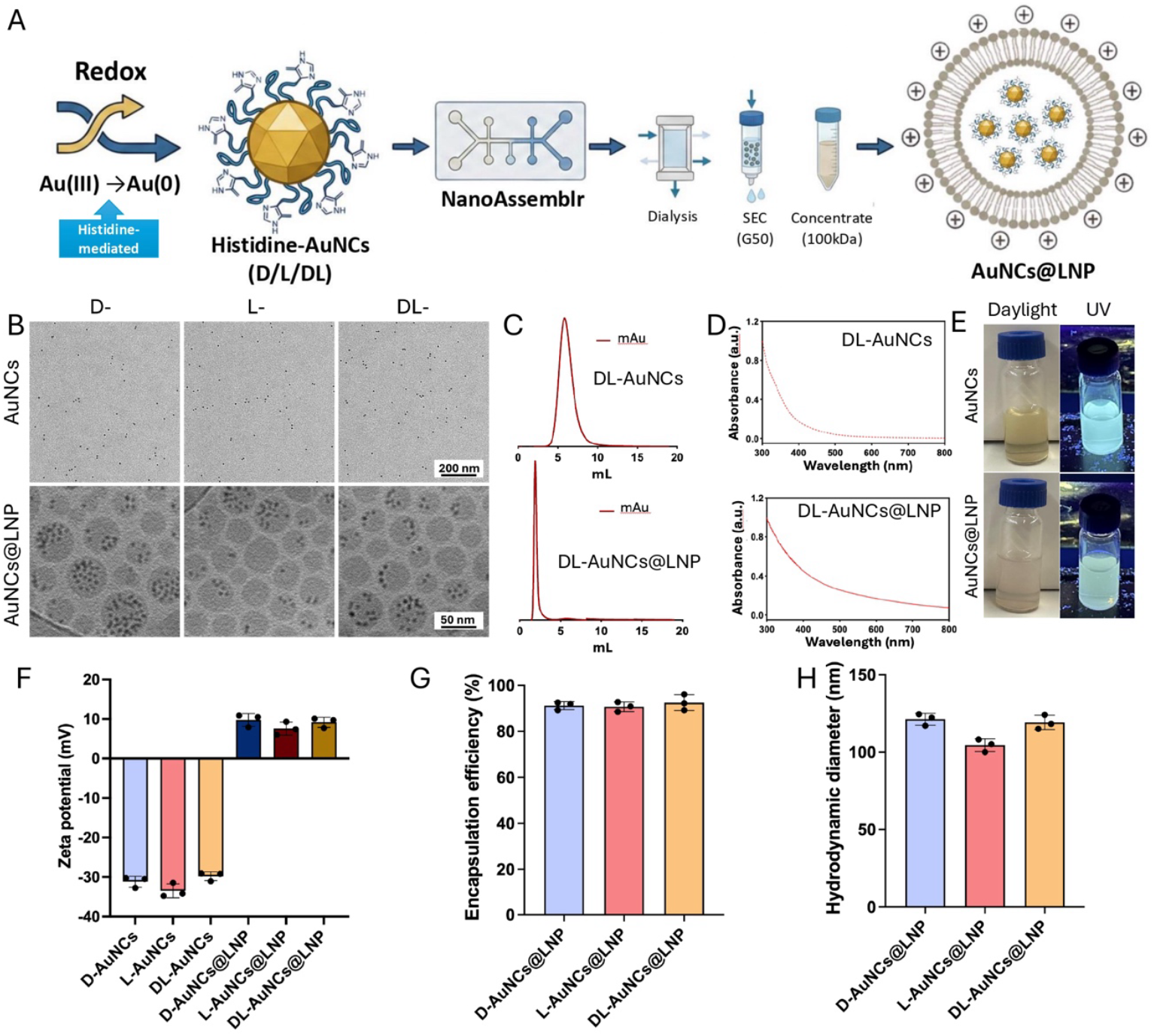
Synthesis, formulation, and physicochemical characterization of chiral AuNCs and AuNCs@LNP. (A) Schematic of His-mediated formation of chiral histidine-stabilized gold nanoclusters, followed by NanoAssemblr microfluidic formulation into cationic lipid nanoparticles and sequential purification (dialysis, size-exclusion chromatography (SEC), and ultrafiltration concentration). (B) Electron microscopy of chiral AuNCs and AuNC@LNPs: TEM images of D-, L-, and DL-AuNCs (top row) showing dispersed ultrasmall electron-dense features without evidence of conversion to larger plasmonic Au nanoparticles within the sampling and resolution of the analysis; cryo-TEM images of the corresponding AuNCs@LNP formulations (bottom row) showing spherical lipid nanoparticles with internal electron-dense inclusions consistent with AuNCs incorporation. Scale bars as indicated. Panels B–E show DL-AuNCs/DL-AuNCs@LNP as representative data; D- and L-variant datasets are provided in the Supporting Information. (C) SEC (Sephadex G-50) elution profiles monitored by mAu (absorbance). Free AuNCs elute later (top). AuNCs@LNPs elute early near the void volume (bottom), indicating separation from unencapsulated AuNCs. (D) UV–vis absorption spectra of AuNCs and AuNCs@LNP showing monotonic absorption without a localized surface plasmon resonance band. (E) Photographs of AuNCs and AuNCs@LNP dispersions under ambient light and 365 nm UV illumination, indicating preserved cyan photoluminescence after LNP formulation. (F) ζ-Potential of free AuNCs (negative) and AuNCs@LNP (mildly positive), supporting installation of a cationic bio-interface. (G) Encapsulation efficiency of AuNCs in LNPs quantified by ICP-OES. (H) Hydrodynamic diameters determined by DLS. Data are mean ± SD (n = 3).

Following microfluidic formulation on the NanoAssemblr platform, cryo-transmission electron microscopy (cryo-TEM) showed predominantly spherical lipid nanoparticles with internal electron-dense inclusions (Figure 1B, bottom), which is consistent with AuNCs incorporation within the LNP interior. To standardize the formulations and, in practical terms, limit residual unencapsulated AuNCs, we applied a sequential purification workflow—dialysis, Sephadex G-50 size-exclusion chromatography, and ultrafiltration—yielding purified AuNCs@LNP dispersions with reduced contributions from free AuNCs (Figure 1C) (36,37). The UV–vis spectra retained the characteristic monotonic absorption profile and lacked a localized surface plasmon resonance band (Figure 1D), supporting preservation of the nanocluster regime rather than conversion into larger AuNCs (38,39). Under the tested conditions, both free AuNCs and AuNCs@LNP remained colloidally stable and continued to exhibit cyan photoluminescence under UV illumination (Figure 1E).

Encapsulation primarily altered interfacial electrostatics, resulting in a charge reversal from negatively charged AuNCs to mildly positive AuNCs@LNP (Figure 1F). This shift supports installation of a cationic interface expected to favor electrostatic interactions with anionic biofilm matrix components (40,41), a design feature that has been implicated as a dominant boundary condition for nanoscale transport and retention in mature biofilms. High encapsulation efficiencies (∼90–94%, quantified by ICP-OES; Figure 1G) together with hydrodynamic diameters of ∼105–122 nm (DLS; Figure 1H) define a modular scaffold in which nanocluster identity is retained while the carrier tunes the biofilm-facing interface, motivating subsequent mechanistic and in vivo evaluation.

### 2.2 Disproportionate biomass and architectural disruption by cationic LNPs relative to CFU reduction

To disentangle bactericidal activity from biofilm matrix/architectural disruption, we compared free AuNCs and AuNCs@LNP against 24 h *S. aureus* USA300 biofilms following 24 h treatment at matched Au doses (25–75 μg/mL Au). Crystal violet (CV) staining showed dose-dependent reductions in retained, adherent biomass for all free AuNC variants (Figure 2A); across the D/L/DL formulations, LNP encapsulation consistently accentuated loss of the adherent fraction (Figure 2A). For example, At 75 μg/mL Au, residual biomass decreased from 100% (PBS) to 63% (DL-AuNCs) and further to 45% with DL-AuNCs@LNP. These data support the view that the mildly cationic lipid shell functions as a matrix-interacting module that promotes detachment and remodels biofilm retention, consistent with reports that positively biased nanocarriers enhance biofilm association and perturb EPS cohesion (42). In parallel, lipid-coated hybrid nanoparticles bearing robust cationic lipid shells have been reported to increase penetration into *S. aureus* USA300 biofilms and improve antibiofilm efficacy relative to non-coated nanoparticles or free antibiotic, underscoring the role of cationic lipid interfaces in governing biofilm association and intrabiofilm transport (31).

**Figure 2.**
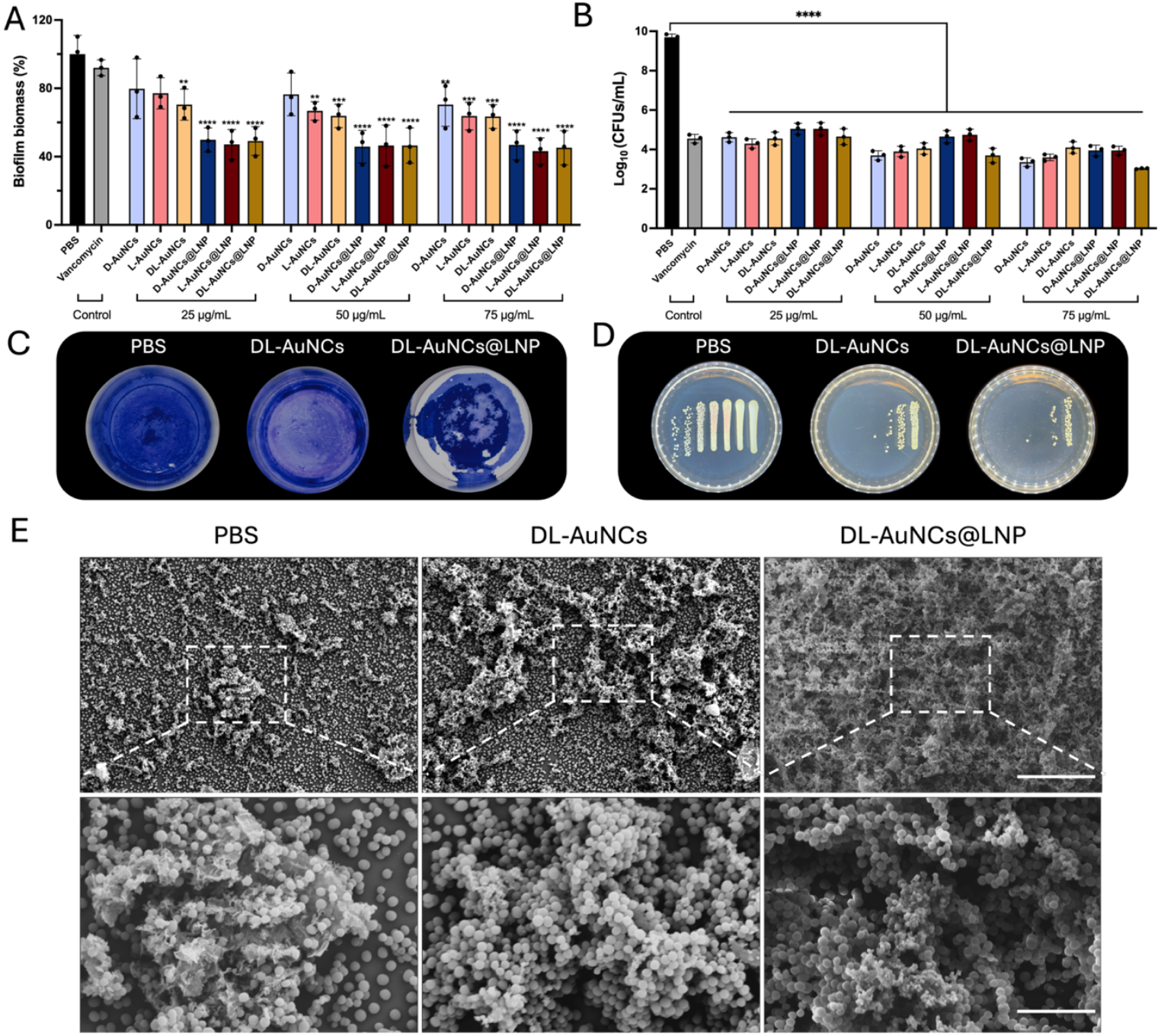
Cationic LNP encapsulation enhances biofilm biomass removal and remodels biofilm structure while maintaining strong bactericidal activity. (A) Residual biofilm biomass of 24 h *S. aureus* USA300 biofilms after 24 h treatment with free AuNCs (D/L/DL) or the corresponding AuNCs@LNP at matched Au doses (25, 50, 75 μg/mLAu), quantified by crystal violet (CV) staining and normalized to PBS control. Vancomycin served as an antibiotic control. (B) Viable bacteria recovered from treated biofilms quantified by CFU enumeration (log_10_ CFU/mL) under the same dosing regimen as in (A). (C) Representative CV-stained wells (left-to-right: PBS, DL-AuNCs, DL-AuNCs@LNP) illustrating residual adherent biomass after the CV workflow (treatment followed by washing and staining). (D) Representative agar plates (left-to-right: PBS, DL-AuNCs, DL-AuNCs@LNP) corresponding to CFU recovery after biofilm disruption and plating. (E) SEM imaging of biofilms directly after the 24 h treatment period (no CV wash/stain workflow), highlighting treatment-dependent differences in biofilm architecture and extracellular matrix–like connectivity (top row, lower magnification; scale bars, 200 μm) and corresponding higher-magnification views (bottom row; scale bars, 5 μm). Data in (A,B) are mean ± SD (n = 3 independent biological replicates; each measured in technical duplicate). A one-way ANOVA with Dunnett’s post-hoc test was used for comparisons against the PBS control group.

In contrast, CFU counting showed robust killing by both free AuNCs and AuNCs@LNP versus PBS across the same dose range (Figure 2B). At 75 μg/mL Au, DL-AuNCs reduced viable burden by 5.6 log_10_, whereas DL-AuNCs@LNP achieved a 6.7 log_10_ reduction, corresponding to an additional ∼1.1 log_10_ gain at matched Au dose. Thus, although LNP formulation can deepen killing under some conditions, its dominant contribution in this mature-biofilm setting is a disproportionate enhancement of biomass/architecture disruption relative to the incremental improvement in CFU reduction. Notably, this decoupling reflects distinct biological readouts rather than an analytical artifact: CV captures retained adherent material (cells plus matrix/debris), whereas CFU quantifies recoverable colony-forming cells after disruption and plating. Accordingly, interventions that weaken matrix cohesion and promote detachment can drive large apparent biomass losses even when additional CFU reductions at a single endpoint are comparatively modest or approach a practical floor (43). The detached small pieces of biofilm have been proved that are more susceptible to the postoperative administrated systemic antibiotics (44).

The SEM dataset provides qualitative support for this interpretation and sets clear limits on what we can claim. In PBS controls, we observed structured coverage with intercellular connectivity consistent with an EPS-rich architecture (Figure 2E). After AuNC treatment, the biofilm surface appeared remodeled. AuNCs@LNP produced a distinct morphology with fewer obvious EPS-like bridges and many small spherical features among cells (Figure 2E). Crystal violet staining did not show an increase in total biomass. This argues against a simple “mass gain” during treatment and suggests that the higher apparent coverage in SEM reflects reorganization and compaction of existing biofilm material, together with fixation/dehydration effects that can collapse hydrated polymer networks. We therefore interpret SEM as evidence of architecture remodeling rather than proof of matrix elimination (45). The small spheres in the AuNCs@LNP condition cannot be assigned unambiguously to intact LNPs or cell-derived debris from SEM alone. Methods such as Au elemental mapping, lipid labeling, or cryo-SEM are needed to resolve particle identity without over-interpreting morphology (45).

### 2.3 Contributions of oxidative stress and metabolic collapse to antibiofilm activity beyond ROS amplitude

To probe mechanism beyond endpoint CFU and biomass, we quantified membrane damage, metabolic activity, and oxidative stress under matched Au dosing. A working model (Figure 3A) is that AuNCs elicit ROS-associated oxidative stress, while cationic LNP encapsulation enhances matrix-level engagement and reshapes how this stress propagates across the biofilm community.

**Figure 3.**
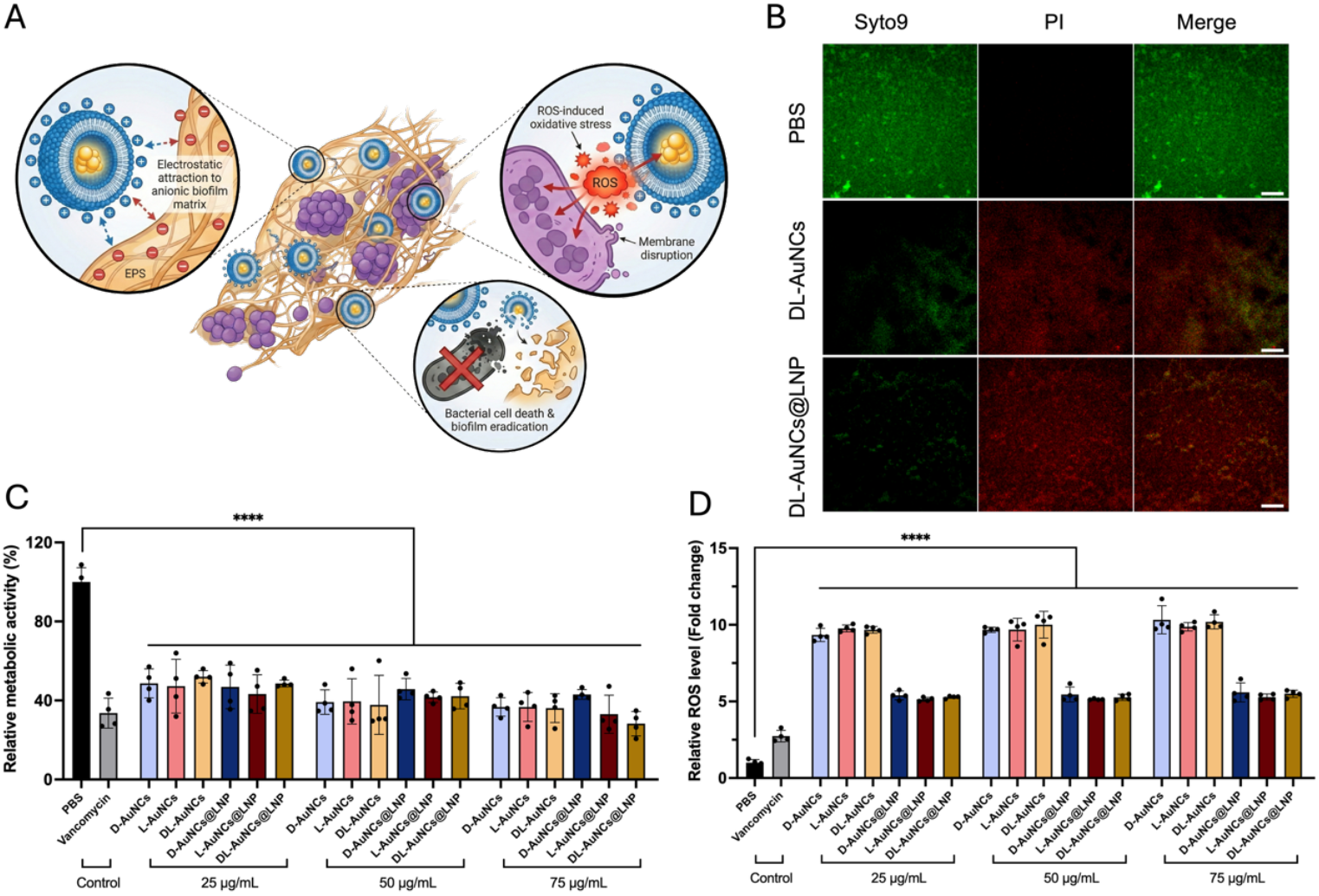
Oxidative stress contributes to AuNC antibiofilm activity, while LNP encapsulation reshapes functional outcomes across orthogonal readouts. (A) Schematic illustrating the proposed sequence of events: electrostatic association of cationic AuNCs@LNP with anionic EPS, local oxidative stress and membrane disruption, and subsequent loss of biofilm viability. (B) Representative Live/Dead CLSM images of 24 h *S. aureus* USA300 biofilms treated for 24 h in PBS with DL-AuNCs or DL-AuNCs@LNP at 75 μg/mL Au (PBS control shown). Green, SYTO 9; red, propidium iodide (PI); merged images shown at right. Scale bars, 10 μm. (C) Biofilm metabolic activity quantified by XTT after 24 h treatment with free AuNCs (D/L/DL) or AuNCs@LNP at matched Au doses (25/50/75 μg/mL); data are expressed as percent of the corresponding PBS control (PBS = 100%). (D) Biofilm-associated ROS measured after 2 h exposure using a fluorescent ROS probe under matched Au dosing (25/50/75 μg/mL); data are plotted as fold change relative to PBS (PBS = 1). DL is shown as representative in panel (B); corresponding D- and L-variant datasets are provided in the Supporting Information. Data in (C,D) are mean ± SD (n = 4 independent biological replicates; each measured in technical duplicate). A one-way ANOVA with Dunnett’s post-hoc test was used for comparisons against the PBS control group.

Live/Dead confocal laser scanning microscopy (CLSM) provided a spatially resolved membrane-integrity readout under a representative condition (DL, 75 μg/mL Au, 24 h treatment). PBS controls were dominated by SYTO 9 signal, whereas DL-AuNCs increased propidium iodide (PI) staining. DL-AuNCs@LNP further intensified PI signal across the field of view (Figure 3B). Because PI reports membrane permeability rather than death per se (46), we interpret these images as an increased fraction of PI-permeable (severely membrane-compromised) cells in the formulated group, consistent with improved biofilm engagement. In parallel, XTT assays across doses (25–75 μg/mL Au) showed that both free AuNCs and AuNCs@LNP suppressed biofilm metabolic activity relative to PBS, with the LNP formulation generally driving a deeper reduction (Figure 3C). As XTT integrates community-level redox activity and respiratory competence rather than culturability per se, this dose-series readout supports an expanded fraction of biofilm-embedded cells experiencing functionally consequential stress upon LNP encapsulation.

ROS measurements add mechanistic context but also define an important interpretive boundary condition. After 2 h exposure, both free AuNCs and AuNCs@LNP elevated ROS above PBS across doses (Figure 3D). Notably, free AuNCs tended to yield higher ROS fold-changes than AuNCs@LNP, even though AuNCs@LNP produced stronger PI staining and more pronounced metabolic suppression (Figure 3B,C). This divergence is plausible in stratified biofilms, where single-time-point fluorescence readouts reflect not only ROS generation but also probe accessibility, local quenching/redox buffering, oxygen gradients, and compartmentalization within EPS(47,48). Accordingly, a lower bulk ROS signal at an early time point does not necessarily indicate a lower time-integrated oxidative burden at the single-cell level. One parsimonious interpretation is that LNP encapsulation redistributes stress across the community, reducing highly localized ROS “hotspots” while increasing the spatial footprint of damaging exposure, thereby producing greater biofilm-wide loss of function at comparable Au dose (47). Alternatively, LNP–matrix and/or LNP– envelope interactions may sensitize cells such that a given ROS increment becomes more lethal. Because we did not perform antioxidant rescue experiments or cell-free probe-interference controls, we interpret the ROS measurements as qualitative support for an oxidative-stress contribution rather than as a standalone quantitative predictor of biofilm killing (48). Collectively, AuNCs elicit ROS-associated stress and metabolic collapse in established *S. aureus* biofilms, while cationic LNP encapsulation shifts how that stress manifests across orthogonal endpoints, enhancing community-level membrane damage and functional suppression even when early bulk ROS amplitude is not maximal.

### 2.4 Favorable in vitro compatibility in cell- and blood-based assays

A translational antibiofilm platform must balance antimicrobial potency with host compatibility, particularly for locally administered formulations. Under the tested conditions, DL-AuNCs and DL-AuNCs@LNP maintained high cell viability in RAW 264.7 macrophages and HeLa cells after 24 h exposure at Au-equivalent concentrations of 50 and 75 μg/mL (Figure 4A,B), with no significant loss relative to untreated controls (P>0.05).These exposure levels align with those used in the antibiofilm assays, supporting an initial cellular compatibility profile relevant to local application scenarios. In addition, hemolysis remained below 5% across the tested Au-equivalent concentrations (Figure 4C), indicating good red blood cell compatibility in line with commonly used hemocompatibility benchmarks (49). In a human whole-blood assay, neither formulation induced measurable increases in IL-6, IL-1β, or TNF-α relative to PBS, whereas LPS produced robust cytokine induction (Figure 4D–F). Together, these data provide an initial safety gate for in vivo evaluation. Nonetheless, standard in vitro assays do not fully capture potential local reactogenicity at concentrated perilesional dosing, nor do they report on complement activation, coagulation/platelet interactions, or tissue-level responses, which remain important considerations for translational development.

**Figure 4.**
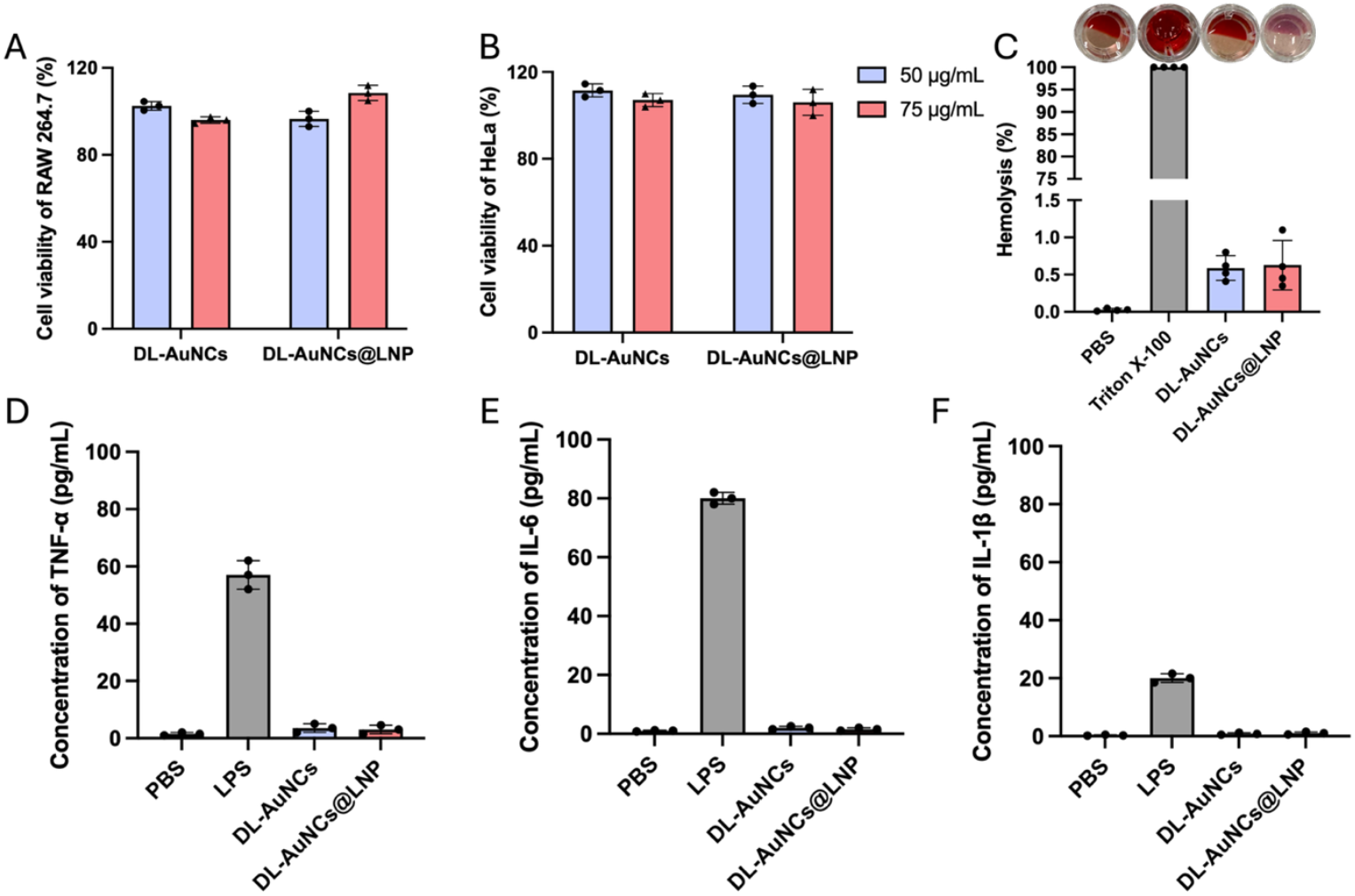
In vitro compatibility of DL-AuNCs and DL-AuNCs@LNP across cellular, blood-cytokine, and hemocompatibility assays. (A,B) Cytocompatibility assessed by MTS in RAW264.7 macrophages (A) and HeLa cells (B) after 24 h exposure to DL-AuNCs or DL-AuNCs@LNP at Au-equivalent concentrations of 50 and 75 μg/mL. Viability is reported relative to untreated controls. (C) Hemolysis of red blood cells incubated with DL-AuNCs or DL-AuNCs@LNP for 1 h; PBS and Triton X-100 serve as negative and positive controls, respectively. (D–F) Human whole-blood cytokine profiling following incubation with DL-AuNCs or DL-AuNCs@LNP, reporting TNF-α (D), IL-6 (E), and IL-1β (F); PBS and LPS serve as negative and positive controls.

### 2.5 In vivo carrier effects and a non-monotonic dose window in implant biofilm therapy

We next asked whether the matrix-engaging carrier advantage observed in vitro translates under the diffusion- and clearance-limited conditions of an implant infection. In a murine subcutaneous catheter biofilm model (Figure 5A), a single perilesional administration of AuNCs@LNP reduced implant-associated bacterial burden relative to PBS and free AuNCs (CFU/implant, day-8 endpoint; Figure 5B). This in vivo separation is consistent with a recurring constraint in implant biofilm therapy: under confined peri-implant delivery, residence and productive interfacial contact at the biofilm–implant boundary can dominate outcome, whereas free nanospecies may be more readily diluted, redistributed, or cleared from the site (50).

**Figure 5.**
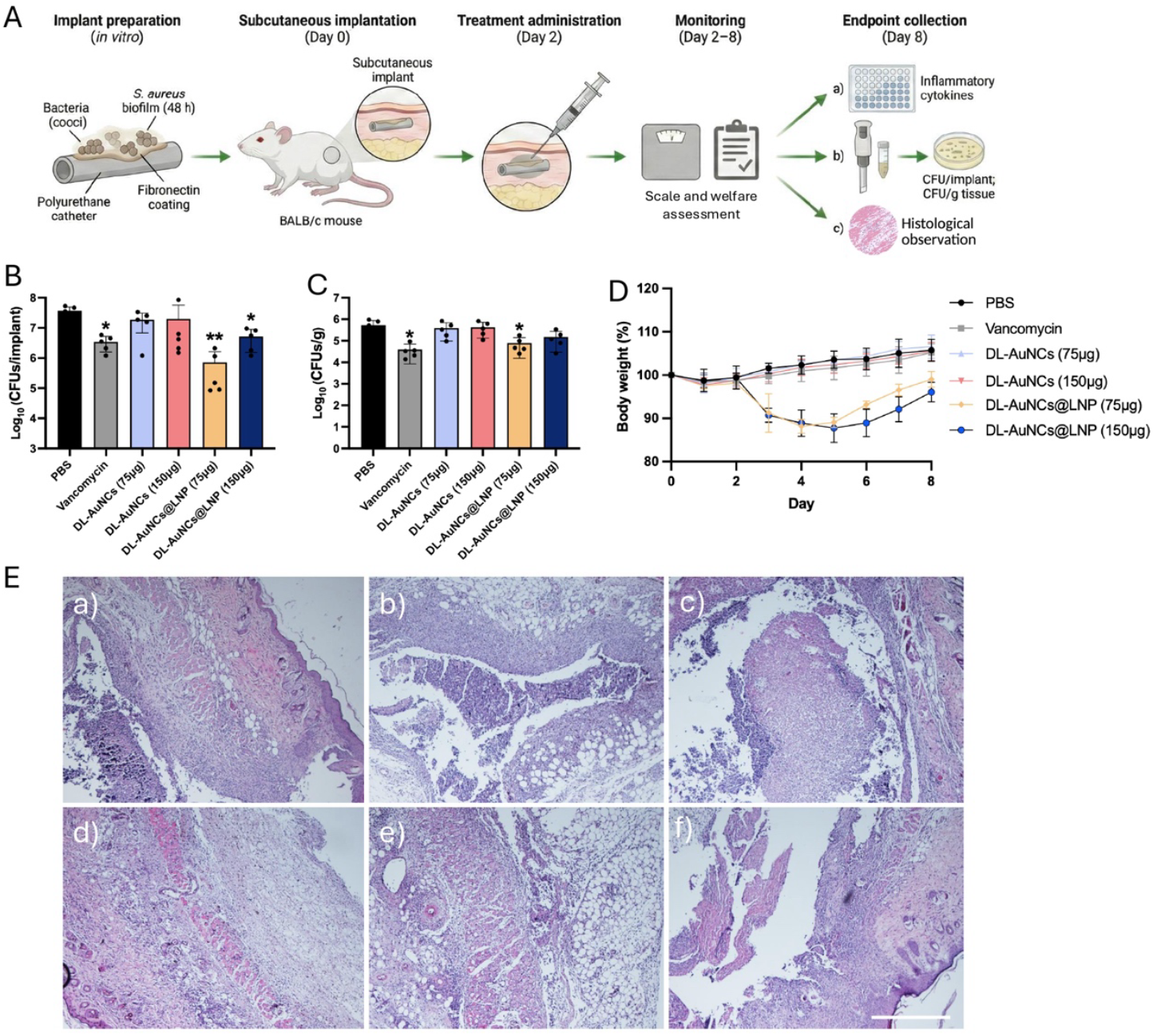
In vivo efficacy and tolerability of AuNCs@LNP in a murine subcutaneous implant–associated *S. aureus* biofilm model. (A) Study schematic. Polyurethane catheters were fibronectin-coated and colonized with *S. aureus* biofilm (48 h) in vitro, implanted subcutaneously in BALB/c mice (day 0), and treated once by perilesional injection on day 2. Animals were monitored (day 2–8) and samples were collected at endpoint (day 8) for inflammatory readouts, bacterial burden, and histology. (B) Implant-associated bacterial burden (CFU/implant) at day 8 following treatment with PBS, AuNCs (75 or 150 μg Au), AuNCs@LNP (75 or 150 μg Au), or vancomycin (500 μg). (C) Peri-implant tissue bacterial burden (CFU/g tissue) at day 8 for the same treatment groups. (D) Body-weight trajectories during the monitoring period (day 2–8) normalized to day-2 baseline. (E) Representative H&E-stained peri-implant tissue sections (day 8) for (E-a) PBS, (E-b) NCs (75 μg Au), (E-c) NCs (150 μg Au), (E-d) NCs@LNP (75 μg Au), (E-e) NCs@LNP (150 μg Au), and (E-f) vancomycin (500 μg). Scale bar, 1 mm. n = 5 mice per group; data are mean ± SD. Statistics: CFU data were log_10_-transformed and analyzed by one-way ANOVA with Bonferroni’s multiple-comparisons test.

A defining feature of these data is the non-monotonic dose response. At matched endpoint, the 75 μg AuNCs@LNP group yielded ∼1.0 log_10_ lower CFU/implant than the 150 μg AuNCs@LNP group (Figure 5B). Peri-implant tissue CFU/g tracked the same overall trend and did not indicate grossly elevated soft-tissue burden under our sampling scheme (Figure 5C), which is compatible with the implant surface remaining the dominant bacterial reservoir and with treatment acting primarily by shaping peri-implant exposure at the biofilm–implant interface rather than broadly sterilizing surrounding tissue. Practically, this is not a simple “more payload, more killing” regime: beyond a threshold, nominal dose increases can become decoupled from productive biofilm exposure as local tissue responses and carrier mass begin to redefine the geometry of delivery.

Tolerability readouts suggest a plausible contributor to this inversion. LNP-containing groups exhibited transient post-injection weight loss and palpable induration, most evident at higher dose (Figure 5D), consistent with a dose- and context-dependent reactogenic response that can remodel the peri-implant microenvironment and thereby reduce productive biofilm contact even when nominal dose increases. Such behavior aligns with prior work showing that strongly cationic lipid nanoparticles can provoke innate immune activation and systemic toxicity markers in vivo (51). In particular, Kedmi et al. compared otherwise similar lipid nanoparticles differing primarily in surface charge and reported that DOTAP-containing positively charged nanoparticles (∼100 nm; ζ ≈ +54 mV) induced body-weight loss and elevated serum liver enzymes after intravenous dosing, accompanied by strong induction of Th1 cytokines and type I interferon–responsive genes in leukocyte populations (52). They further showed that this pro-inflammatory program was largely TLR4-dependent and could be triggered directly at the leukocyte level, underscoring how positively biased lipid surfaces can impose dose ceilings through host sensing pathways. Although our AuNCs@LNP are only mildly cationic and are delivered locally rather than systemically, this literature supports the broader interpretation that increasing lipid dose can amplify host–material interactions in a non-linear fashion and thereby reshape the effective exposure geometry in complex tissues.

Histology (H&E) showed inflammatory remodeling across infected groups, as expected in an implant bed. AuNCs@LNP groups displayed prominent clear spaces within reactive regions (Figure 5E-d,e). We interpret these features cautiously, for example as lipid-associated vacuolation versus processing-related lipid extraction. However, their enrichment in the higher-dose LNP condition, together with systemic and local signs, supports the broader conclusion that host–material interactions can impose a ceiling on effective local dosing.

From a spatial-pharmacology perspective, a parsimonious model is that increasing LNP dose moves the system from a coverage-limited regime, where added particles increase implant-adjacent coverage and matrix engagement, to a host-limited regime, where added lipid predominantly amplifies local reactogenic remodeling and/or clearance rather than increasing productive biofilm contact. Three non-exclusive mechanisms can account for this transition, each testable without overcommitting interpretation. First, depot-like confinement or inflammatory encapsulation may reduce circumferential distribution around the implant. Second, phagocyte-mediated sequestration and clearance may accelerate in a more inflamed niche. Third, microaggregation and protein adsorption at high local concentration may reduce the usable interfacial particle population (53). Notably, all three are consistent with the key empirical constraint here. Higher-dose AuNCs@LNP did not improve CFU/implant despite delivering more Au, which implies that effective biofilm exposure is limited by local biology and carrier behavior rather than intrinsic antimicrobial capacity.

Practically, optimization should focus on widening the efficacy–tolerability window. For example, this may be achieved by moderating effective cationic surface density while preserving matrix association, tuning lipid composition and PEG-lipid content to reduce reactogenicity, or implementing split dosing to sustain exposure without exceeding local tolerance. The most direct next experiments to close the loop are (1) circumference- and depth-resolved peri-implant distribution mapping (fluorescence plus elemental Au), (2) time-resolved clearance with immune colocalization, and (3) histology with lipid and immune markers to distinguish lipid-associated tissue reactions from processing artifacts (50).

## 3. Conclusion

In summary, we developed a modular antibiofilm nanomedicine in which chiral histidine-stabilized gold nanoclusters provide a bactericidal component associated with physiological stress, while cationic lipid nanoparticle encapsulation reconfigures the biofilm-facing interface to promote matrix-level engagement without conversion of AuNCs into plasmonic gold nanoparticles. Across established *S. aureus* USA300 biofilms, orthogonal readouts indicated a partial decoupling between killing and biomass/architecture outcomes: AuNCs drove pronounced reductions in recoverable viability, whereas AuNCs@LNP disproportionately enhanced loss of retained adherent biomass and remodeled biofilm ultrastructure, consistent with carrier-mediated perturbation of matrix cohesion and/or detachment. Mechanistic assays further supported community-level metabolic suppression and ROS-associated stress, while underscoring that early bulk ROS amplitude is not a monotonic predictor of lethality in stratified biofilms.

The platform showed favorable in vitro compatibility across cellular viability and hemocompatibility assays. In vivo, AuNCs@LNP improved implant-associated burden in a murine subcutaneous catheter biofilm model, but efficacy followed a non-monotonic dose window in which higher carrier exposure coincided with tolerability constraints consistent with local reactogenic responses. Together, these findings provide an early preclinical proof-of-concept that AuNCs@LNP can couple nanocluster-mediated antibacterial activity with lipid-facilitated biofilm disruption, while highlighting exposure- and host-limited constraints that are particularly relevant to local delivery. Nevertheless, our in vivo evidence is currently restricted to a subcutaneous catheter model and does not capture the anatomical, mechanical, or microbiological complexity of orthopedic implant-associated bone and joint infections. Translation toward orthopedic indications will therefore require validation in load-bearing bone/joint implant models and evaluation alongside standard surgical debridement and antibiotic regimens before clinical translation can be considered.

## 4. Methods

### 4.1 Bacterial Strain and Growth Conditions

*Staphylococcus aureus* USA300 LAC (AH1263) strain was used in this study (54). The strain was streaked onto sheep blood agar (SBA) plates and incubated overnight at 37 °C. A single colony was inoculated into tryptic soy broth (TSB) and cultured overnight at 37 °C with shaking (600 rpm). For each experiment, the overnight culture was diluted into fresh TSB and grown for 3 h at 37 °C to obtain logarithmic-phase cells.

### 4.2 Synthesis of Chiral Gold Nanoclusters

Chiral histidine-functionalized gold nanoclusters were prepared using a published method with minor modifications (24,25). Briefly, HAuCl_4_ (10 mM, 1.0 mL; Sigma-Aldrich) was mixed with D-, L-, or DL-histidine (100 mM, 3.0 mL; Sigma-Aldrich) at room temperature under stirring (600 rpm). The resulting products are referred to as D-AuNCs, L-AuNCs, and DL-AuNCs, respectively. The reaction proceeded for 16 h in the dark at room temperature. The mixture was passed through a 0.2 µm syringe filter and concentrated using 3 kDa Amicon Ultra centrifugal filters (Millipore) to the desired Au concentration. Au concentration was quantified by ICP-OES (see below). AuNCs were stored at 4 °C protected from light.

### 4.3 Preparation of Chiral AuNC-Loaded Lipid Nanoparticles

AuNCs@LNPs were engineered via an optimized self-assembly process based on the foundational lipid-ionizable frameworks developed by Zhigaltsev et al (55). LNPs were formulated using DLin-MC3DMA/DSPC/cholesterol/PEG-DMG at a molar ratio of 50.0/10.0/38.5/2.0. Lipids were dissolved in ethanol at a total lipid concentration of 10 mM. In parallel, AuNCs were prepared in 25 mM sodium acetate buffer (pH 4.0) at an Au concentration of 0.2 mg/mL. Nanoparticles were assembled using a NanoAssemblr microfluidic mixer (Precision NanoSystems) by combining ethanol and aqueous streams at a 1:3 (v/v) ratio with a total flow rate of 9 mL/min at room temperature. The resulting suspension was dialyzed against PBS (pH 7.4) using a 10 kDa MWCO dialysis cassette at 4 °C for ∼16 h with three buffer exchanges. Formulations were purified by Sephadex G-50 (Cytiva, UK) size-exclusion chromatography to remove unencapsulated AuNCs (column bed volume ∼10 mL; 0.5 mL fractions collected; nanoparticle-containing fractions identified by turbidity and pooled). Purified AuNCs@LNPs were concentrated to the desired Au concentration using 100 kDa Amicon Ultra filters and stored at 4 °C.

### 4.4 Characterization of AuNCs and AuNCs@LNPs

#### Hydrodynamic Diameter and Polydispersity Index

Hydrodynamic diameter, polydispersity index (PDI), and size distribution were measured by dynamic light scattering (DLS; Malvern Panalytical). Samples were diluted 1:20 in PBS and measured at 25 °C using disposable polystyrene cuvettes. Each formulation was measured in triplicate with 20 runs per measurement.

#### Zeta Potential

Zeta potential was determined by laser Doppler electrophoresis using a Zetasizer Nano Z (Malvern Panalytical). Samples were diluted in 20 mM HEPES buffer (pH 7.4) and measured in three independent runs.

#### UV–Vis Absorption Spectroscopy

UV–vis spectra were collected on a Shimadzu UV-2450 spectrophotometer over 300–800 nm. Samples were diluted in PBS prior to measurement.

#### Gold Element Concentration and Encapsulation Efficiency

Gold concentrations in AuNCs and AuNCs@LNPs were determined by inductively coupled plasma optical emission spectroscopy (ICP-OES; Avio 500, PerkinElmer, USA) using an Au standard curve prepared in deionized water. Samples were digested in freshly prepared aqua regia (HCl/HNO_3_, 3:1, v/v) for 30 min (or until complete digestion) and diluted with deionized water prior to analysis. Au was quantified using the instrument Au emission line at ∼267 nm. Encapsulation efficiency (EE%) was calculated as:

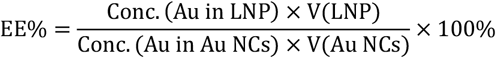

#### Transmission electron microscopy (TEM) and cryogenic transmission electron microscopy (Cryo-TEM)

For TEM, 10 µL of AuNC solution was applied to Quantifoil copper grids, blotted, and air-dried overnight. Grids were imaged using a transmission electron microscope (FEI Tecnai series). Particle features were quantified in Fiji. For cryo-TEM, 10 µL of AuNCs@LNP suspension was applied to freshly glow-discharged Quantifoil grids in a humidified chamber, blotted, and vitrified using an FEI Vitrobot Mark IV (blot time 3 s; blot force 0; humidity 100%). Grids were stored in liquid nitrogen and imaged on an FEI Tecnai G2 20 TWIN microscope operated at 200 kV with a Gatan 70° tilt cryo-transfer system. Micrographs were collected at 29,000× magnification using an FEI 4k × 4k Eagle camera.

### 4.5 In Vitro Biofilm Assays

#### Biofilm Culture

Overnight cultures were adjusted to OD600 = 1.0 in TSB supplemented with 0.5% (w/v) glucose and 3% (w/v) NaCl, then diluted 1:1000 to an initial inoculum of ∼10^5^ CFU/mL. Aliquots (200 µL) were added to tissue-culture treated 96-well plates (Corning) and incubated statically for 24 h at 37 °C. For enhanced adhesion, plates were coated with bovine fibronectin (20 µg/mL in 0.1 M carbonate–bicarbonate buffer, pH 9.4–9.8) at 4 °C overnight, followed by one PBS wash prior to seeding.

#### Crystal Violet Biofilm Biomass Quantification

Biofilms were gently washed with PBS to remove planktonic cells and treated with D-, L-, or DL-AuNCs with or without LNP formulation at matched Au doses (25, 50, and 75 µg/mL Au) for 24 h at 37 °C. PBS served as a negative control and vancomycin (250 µg/mL) as a positive control. After treatment, biofilms were washed with PBS, heat-fixed at 60 °C for 60 min, stained with 0.1% (w/v) crystal violet for 5 min, and rinsed thoroughly with water. After drying, bound dye was eluted with 33% (v/v) acetic acid and shaken for 5 min. Absorbance was measured at 595 nm.

#### Colony-Forming Unit (CFU) Enumeration

After 24 h biofilm formation and treatment as above, biofilms were washed with PBS and detached by sonication in PBS for 5 min using a bath sonicator (40 kHz, 180 W; on ice), followed by vigorous pipetting. Serial 10-fold dilutions (10^−1^ to 10^−6^) were prepared in sterile PBS, and 2 µL spots were plated onto TSA. Plates were incubated at 37 °C for 18–24 h prior to colony enumeration.

#### XTT Metabolic Activity Assay

After treatment, biofilms were gently washed with PBS. XTT reagent (Biotium) was prepared by mixing 25 µL activation reagent with 5 mL XTT solution. XTT (25 µL per well) was added directly to the biofilm-containing wells and incubated at 37 °C in the dark for 2 h. Absorbance was measured at 490 nm with background subtraction. Data were normalized to PBS controls.

#### Reactive Oxygen Species Assay

Biofilms were stained with 1× ROS probe solution (prepared from a 500× stock in DMSO) for 60 min at 37 °C in the dark. Without washing, AuNCs or AuNCs@LNPs were added at 25, 50, and 75 µg/mL Au. Controls included PBS (negative control), H_2_O_2_ (100 µM; positive control), and vancomycin (250 µg/mL and 62.5 µg/mL, ± LNP). ROS fluorescence was measured on a microplate reader (Ex 495 nm/Em 520 nm) at a fixed endpoint of 2 h. Fluorescence values were normalized to biofilm biomass determined by crystal violet staining at 595 nm in parallel identically treated wells. Experiments were performed in quadruplicate under dark conditions.

#### Live/Dead Confocal Microscopy

Biofilms were grown on sterile glass disks placed in 24-well plates by inoculating ∼10^6^ CFU/mL bacteria and incubating for 24 h at 37 °C. After washing three times with 0.85% NaCl, biofilms were treated with AuNCs@LNP (75 µg/mL Au) for 24 h. Biofilms were stained using the LIVE/DEAD BacLight kit (Thermo Fisher) with SYTO 9 and propidium iodide at final concentrations of 6 µM and 30 µM, respectively, for 15 min in the dark. Disks were rinsed gently, mounted on slides, and imaged by CLSM using a 20× objective. Z-stacks were acquired with a step size of 2 µm across the full biofilm thickness. Fluorescence channels were collected at 488 nm excitation (SYTO 9; 520/40 nm) and 535 nm excitation (PI; 560/40 nm).

#### Scanning Electron Microscopy (SEM)

Biofilms were grown on poly-L-lysine-coated 12 mm glass coverslips and treated for 6 h, 12 h, 24 h at 75 µg/mL Au. Samples were fixed for 24 h at room temperature in 2% formaldehyde, 0.5% glutaraldehyde, and 0.15% ruthenium red in 0.1 M phosphate buffer (pH 7.4), rinsed twice, and post-fixed for 2 h at 4 °C in 1% osmium tetroxide and 1.5% potassium ferricyanide in 0.065 M phosphate buffer. After dehydration in graded ethanol (50%, 70%, 80%, 95%, 2×100%), samples were exchanged into HMDS (50% HMDS/ethanol, 2×100% HMDS) and air-dried overnight. Samples were sputter-coated with 6 nm Au (Quorum Q150R S) and imaged on a Zeiss Gemini 450 at 3 kV with a working distance of 4–6 mm.

### 4.6 In Vitro Cytotoxicity and Immunogenicity Assays

#### Cell Viability (MTS)

HeLa cells were cultured in DMEM supplemented with 10% fetal bovine serum (FBS) and 1% penicillin/streptomycin at 37 °C with 5% CO_2_. RAW264.7 murine macrophages were maintained under the same conditions. Cells were seeded into 96-well plates (HeLa: 5,000 cells/well; RAW264.7: 10,000 cells/well) and allowed to adhere overnight prior to treatment. AuNCs or AuNCs@LNPs were added at 50 and 75 µg/mL Au and incubated for 24 h. Cell viability was assessed using the CellTiter 96 AQueous One Solution Cell Proliferation (MTS) assay (Promega) according to the manufacturer’s instructions. Briefly, 20 µL of MTS reagent was added to each well containing 100 µL medium and incubated for 1–2 h at 37 °C protected from light. Absorbance was recorded at 490 nm using a SpectraMax ID3 plate reader (Molecular Devices). Untreated cells were set to 100% viability, and background absorbance (medium + MTS without cells) was subtracted.

#### Whole Blood Cytokine Assay

Peripheral whole blood was collected from healthy donors after informed consent and diluted 1:5 in plain RPMI (without FBS or antibiotics). Diluted blood (600 µL) was dispensed into 48-well plates and exposed to AuNCs or AuNCs@LNPs at final Au concentrations of 25 or 50 µg/mL, with RPMI added as needed to keep the total volume constant. RPMI alone served as the negative control. Lipopolysaccharide (LPS, 1 ng/mL) was used as a positive control for cytokine induction, and PMA (25 ng/mL) served as a positive control for oxidative burst activity. Following incubation for 18–24 h at 37 °C and 5% CO_2_, samples were centrifuged at 500 × g for 5 min. Supernatants were harvested and stored at −80 °C until measurement. IL-6, TNF-α, and IL-1β were quantified using DuoSet ELISA kits (R&D Systems) according to the manufacturer’s protocol. Absorbance was recorded at 450 nm with background correction at 540 nm.

#### Hemolysis Assay

Fresh human red blood cells were isolated from whole blood by centrifugation (800 × g, 5 min) and washed three times with PBS. RBCs were resuspended to 2% (v/v) in PBS and incubated with AuNCs or AuNCs@LNPs at 25 and 50 µg/mL Au for 1 h at 37 °C. PBS served as a negative control, and Triton X-100 (0.1% v/v) served as a positive control. Samples were centrifuged (800 × g, 5 min) and supernatant absorbance was measured at 540 nm. Hemolysis (%) was calculated relative to Triton X-100.

#### Ethics Statement

Human blood was obtained from healthy donors who provided informed consent. Procedures complied with the Declaration of Helsinki and were approved by the Medical Ethics Committee of the University Medical Center Utrecht (METC protocol 07-125/C; approved March 1, 2010).

### 4.7 In Vivo Murine Subcutaneous Biofilm Implant Model

The murine subcutaneous implant-associated biofilm infection model was performed according to a previously described protocol with minor modifications (56). Male Balb/cAnNCrl mice (Charles River Laboratories, Germany; >20 g, 7–10 weeks old) were housed under standard conditions with ad libitum access to food and water. All procedures were approved by the Animal Welfare Body Utrecht (AWB Utrecht) under project license AVD11500202518775 and complied with Directive 2010/63/EU. Seven French polyurethane catheters were sterilized with 70% ethanol, air-dried, and coated with bovine fibronectin overnight at 4 °C. Catheters were incubated in ∼10^7^ CFU/mL *S. aureus* in TSB for 48 h at 37 °C with shaking (250 rpm), washed three times with PBS to remove planktonic bacteria, and stored in PBS until implantation. Mice were acclimated for 7 days prior to surgery. Anesthesia was induced with isoflurane (5%) and maintained at 2% in oxygen, and ocular lubricant was applied. Buprenorphine (0.075 mg/kg, s.c.) was administered at least 30 min preoperatively. After shaving and disinfection, a 5 mm incision was made in the interscapular region, a 14G needle was inserted subcutaneously, and a 5 mm colonized catheter segment was positioned using a K-wire. The incision was closed with 5-0 absorbable sutures, and animals recovered on a 37 °C warming pad; buprenorphine was continued every 12 h for 48 h. Animals were randomized into six groups (n = 5 per group). On day 2 post-implantation, a single 200 µL subcutaneous injection was administered around the infection site using a 29G insulin syringe. Treatments were PBS; AuNCs (75 µg or 150 µg Au per mouse); AuNCs@LNP (75 µg or 150 µg Au per mouse); and vancomycin (500 µg per mouse). Personnel performing injections and outcome analyses were blinded to group assignment. Animals were monitored daily and body weight was recorded each day. Welfare was assessed using a cumulative scoring system evaluating coat condition, posture, activity level, weight loss, implant-site appearance, respiratory status, and overall condition. Humane endpoints were defined as a total welfare score ≥15 or ≥20% body weight loss, at which point animals were euthanized. On day 8, mice were euthanized by CO_2_ followed by cervical dislocation. Blood was collected by cardiac puncture; 20 µL whole blood was plated directly for CFU, and the remaining blood was processed to serum (3,000 × g, 10 min, 4 °C). Implants and peri-implant tissue were harvested aseptically. Tissue was homogenized in PBS using a bead homogenizer (30 Hz, 2 min), serially diluted, and plated on TSA. Plates were incubated at 37 °C for 18–24 h before CFU counting. Bacterial burden was expressed as CFU/implant and CFU/g tissue. Peri-implant tissues for histology were fixed in 10% neutral buffered formalin for 24 h, transferred to 70% ethanol, embedded in paraffin, sectioned at 5 µm, and stained with hematoxylin and eosin (H&E). Sections were imaged on an Olympus BX51 microscope.

### 4.8 Statistical Analysis

All analyses were performed in GraphPad Prism 10. In vitro datasets were analyzed using one-way ANOVA followed by Bonferroni’s multiple-comparisons test. In vivo CFU/implant data were log_10_-transformed and analyzed by one-way ANOVA with Bonferroni post hoc correction. Unless stated otherwise, data are presented as mean ± SD. In vitro experiments were performed with n ≥ 3 independent biological replicates, and in vivo experiments used n = 5 per group with blinded administration and analysis. A two-sided p < 0.05 was considered statistically significant.

## Abbreviations

AMR: antimicrobial resistance
ANOVA: analysis of variance
AuNCs: gold nanoclusters
AWB: Animal Welfare Body
BSA: bovine serum albumin
CFU: colony-forming unit
CLSM: confocal laser scanning microscopy
Cryo-TEM: cryogenic transmission electron microscopy
DLS: dynamic light scattering
DMSO: dimethyl sulfoxide
EE: encapsulation efficiency
ELISA: enzyme-linked immunosorbent assay
EPS: extracellular polymeric substance
FBS: fetal bovine serum
GDL: Gemeenschappelijke Dieren Laboratorium
H&E: hematoxylin and eosin
HEPES: 4-(2-hydroxyethyl)-1-piperazineethanesulfonic acid
AuNCs@LNP: histidine-modified gold nanocluster-loaded lipid nanoparticles
HMDS: hexamethyldisilazane
HRP: horseradish peroxidase
ICP-OES: inductively coupled plasma optical emission spectroscopy
IL-1β: interleukin-1 beta
IL-6: interleukin-6
LNPs: lipid nanoparticles
LPS: lipopolysaccharide
MBBC: minimum biofilm bactericidal concentration
METC: Medical Ethics Committee: *Staphylococcus aureus* (*S. aureus*)
MRSA: methicillin-resistant *Staphylococcus aureus*
MTS: 3-(4,5-dimethylthiazol-2-yl)-5-(3-carboxymethoxyphenyl)-2-(4-sulfophenyl)-2H-tetrazolium
NCs: nanoclusters
OD: optical density
PBS: phosphate buffered saline
PDI: polydispersity index
PEG-DMG: polyethylene glycol-dimyristoyl glycerol
PI: propidium iodide
PMA: phorbol 12-myristate 13-acetate
ROS: reactive oxygen species
RPMI: Roswell Park Memorial Institute medium
SBA: sheep blood agar
SEM: scanning electron microscopy
TEM: transmission electron microscopy
TNF-α: tumor necrosis factor-alpha
TSA: tryptic soy agar
TSB: tryptone soy broth
UV-Vis: ultraviolet-visible
WHO: World Health Organization
XTT: 2,3-bis-(2-methoxy-4-nitro-5-sulfophenyl)-2H-tetrazolium-5-carboxanilide

## Acknowledgments

This publication is part of the project DARTBAC (with project number NWA.1292.19.354) of the research program NWA-ORC which is (partly) financed by the Dutch Research Council (NWO). This work was supported by the European Union’s Horizon 2020 research and innovation program [Grant Agreement No. 825828]. We thank Corlinda ten Brink, Tineke Veenendaal, Cilia de Heus and the Cell Microscopy Core team, UMCU, for technical support. We thank Eric Hellebrand, Schneijdenberg, Erik Betz-Güttner and the Electron Microscopy Centre at Utrecht University, for technical support. We thank Helen de Waard, Afshin Neshad Ashkzar and ICP Lab at Utrecht University. We also thank Dr. Tim Sakkers, Dr. Jort van der Geest, and Dr. Michele Buono, Dr. Qiangbin Yang, Dr. Geng Yang for their support. Generative AI tools (Gemini, ChatGPT, Grok) were used to assist with grammar correction and improving clarity. The authors take full responsibility for the content and accuracy of the paper.

