## Supplementary Information for "Chiral Histidine-Modified Gold Nanoclusters Loaded into Cationic Lipid Nanoparticles for Treatment of Biofilm- associated Infections"

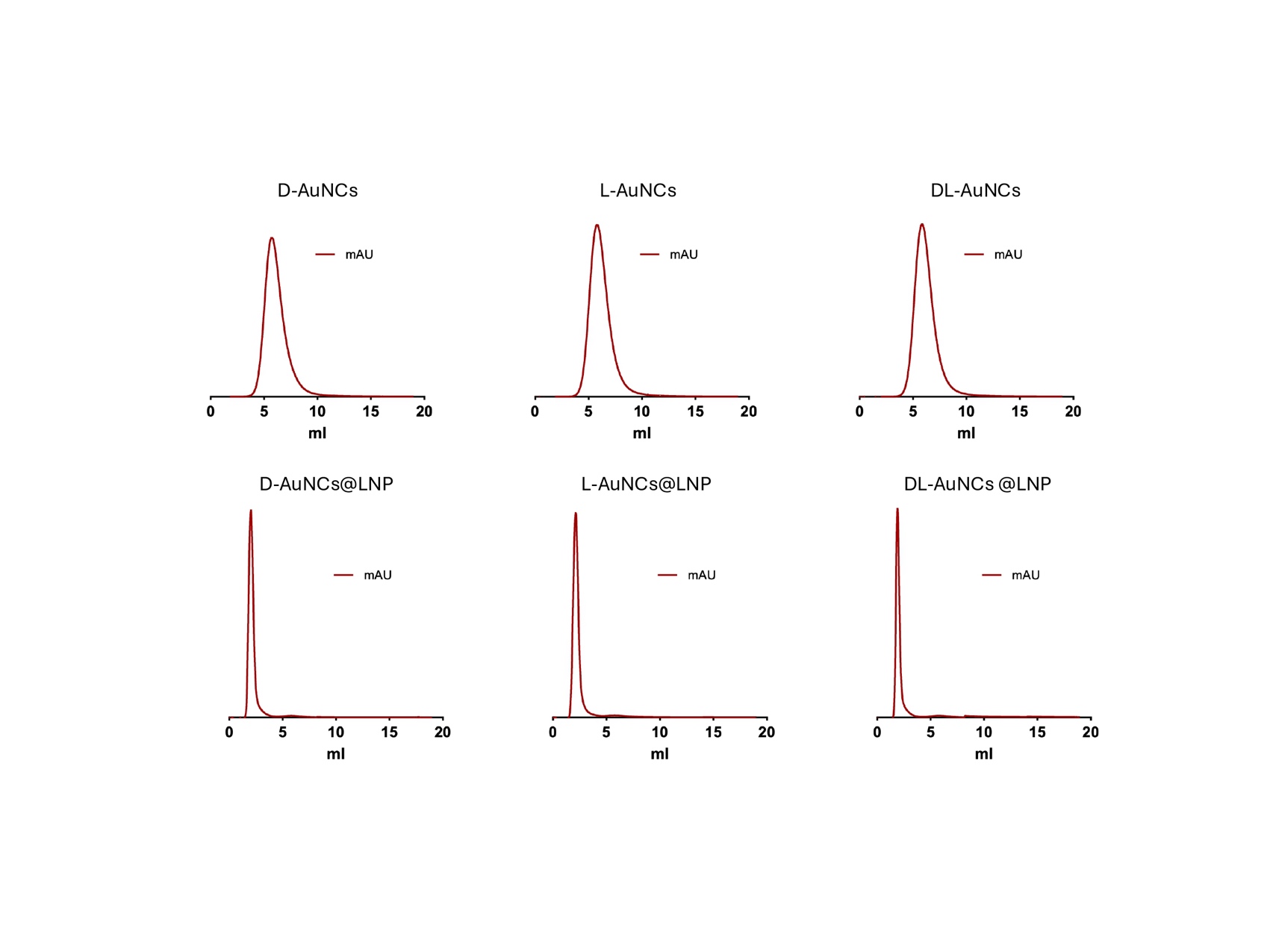

**Figure S1. Size-exclusion chromatography (SEC) elution profiles of chiral AuNCs and AuNCs@LNP.** Representative SEC chromatograms (UV absorbance, mAU, plotted versus elution volume, mL) for D-, L-, and DL-AuNCs (top row) and the corresponding D-, L-, and DL-AuNCs@LNP formulations (bottom row). Free AuNCs elute at higher volumes, whereas AuNCs@LNP elute near the column void volume, consistent with a larger hydrodynamic size and effective separation of nanoparticle-associated material from unencapsulated AuNCs.

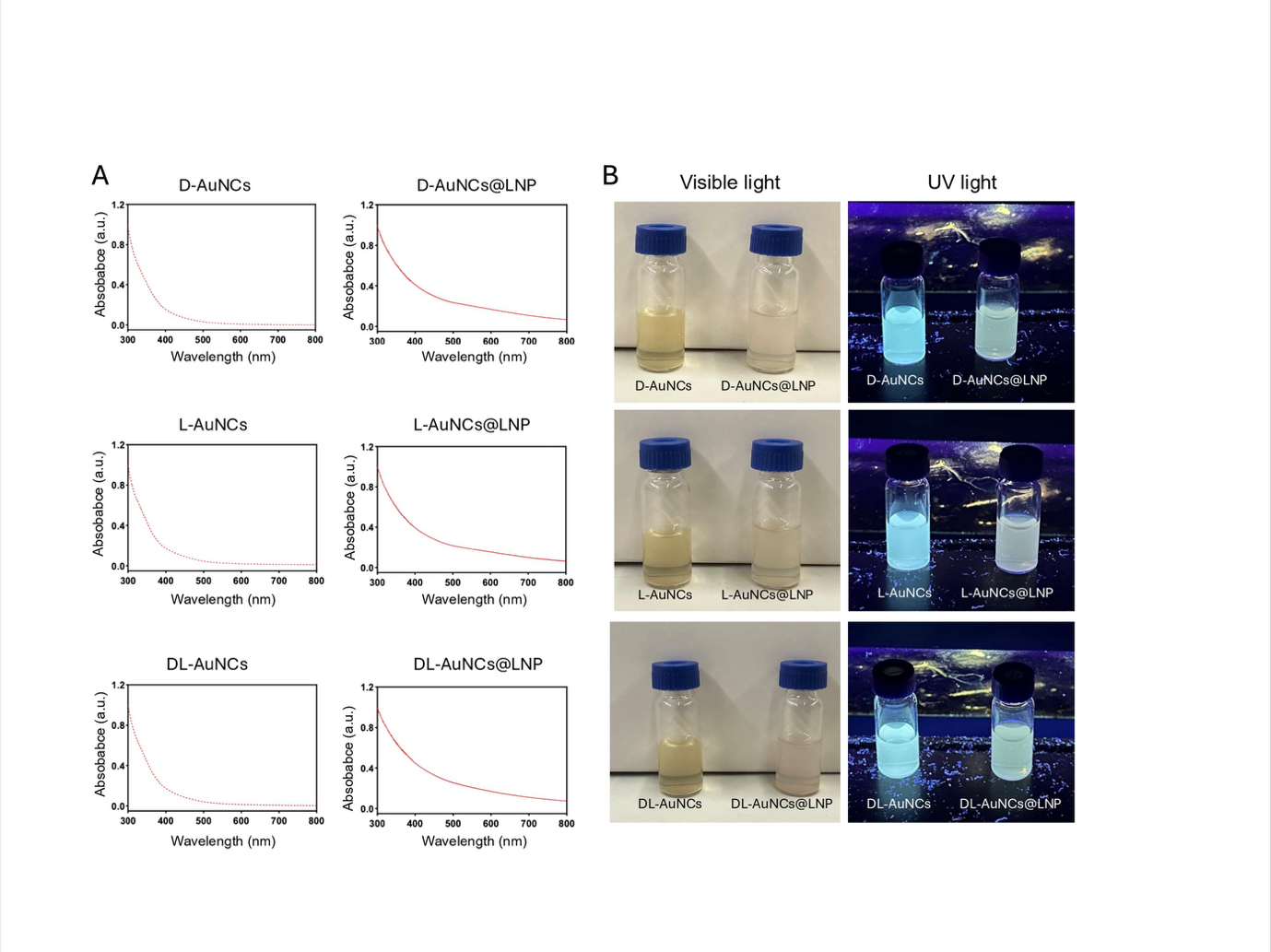

**Figure S2. UV–vis absorption and photoluminescence of chiral AuNCs before and after LNP encapsulation.** (A) UV–vis spectra (300–800 nm) of D-, L-, and DL-AuNCs and the corresponding AuNCs@LNP formulations. All samples show the characteristic monotonic nanocluster absorption profile and lack a localized surface plasmon resonance band, consistent with preservation of the nanocluster regime after formulation. (B) Representative photographs of the same formulations under visible light and 365 nm UV illumination, showing retained cyan photoluminescence following LNP encapsulation.

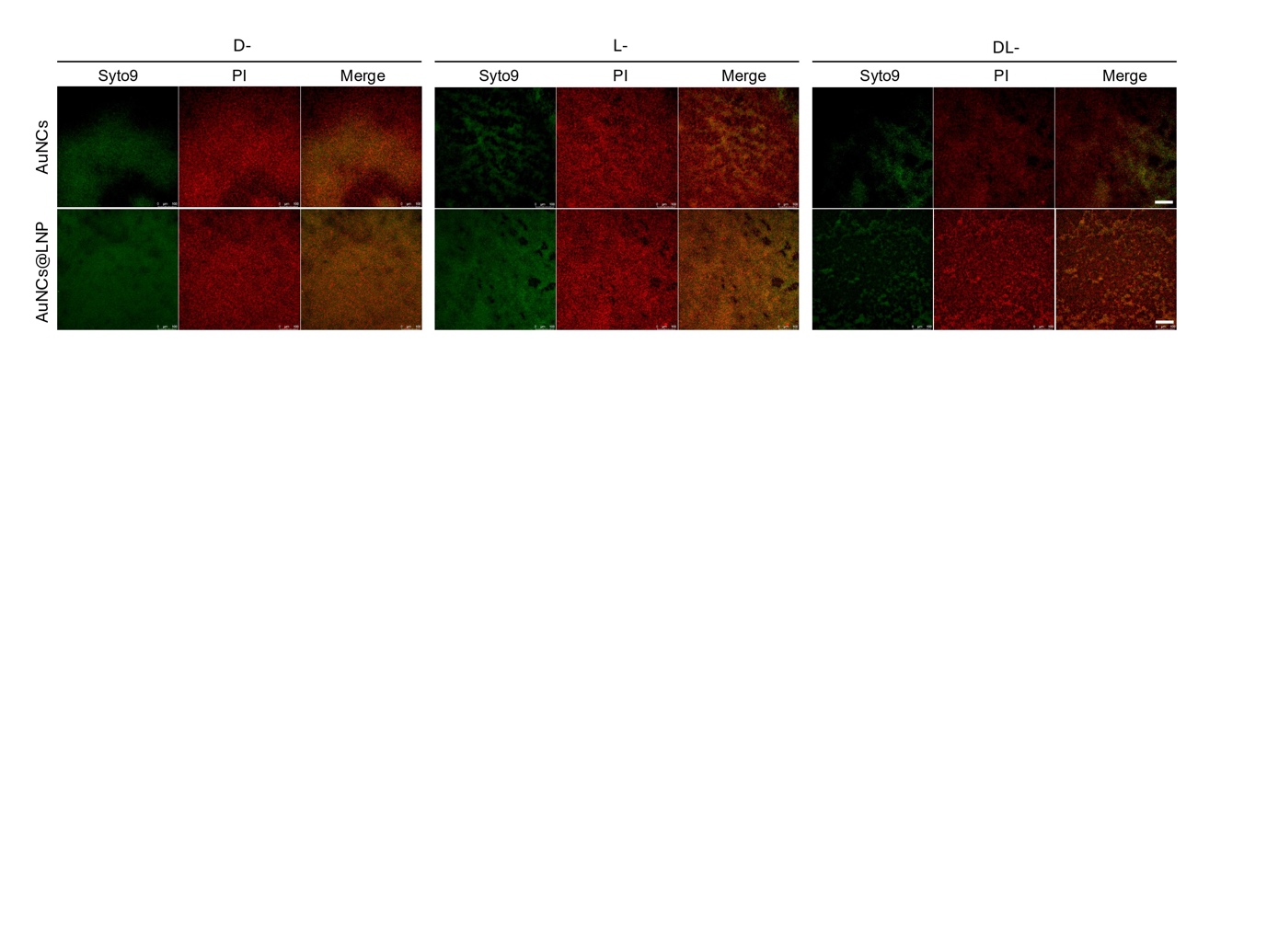

**Figure S3. Live/Dead CLSM imaging of *S. aureus* USA300 biofilms treated with chiral AuNCs or AuNCs@LNP at 75 μg/mL Au.** Representative confocal micrographs of 24 h biofilms after 24 h exposure to D-, L-, or DL-AuNCs (top row) or the corresponding AuNCs@LNP formulations (bottom row) at an Au-equivalent concentration of 75 μg/mL. Membrane integrity was assessed using SYTO 9 (green) and propidium iodide (PI; red); merged channels are shown. Scale bars, 10 μm.

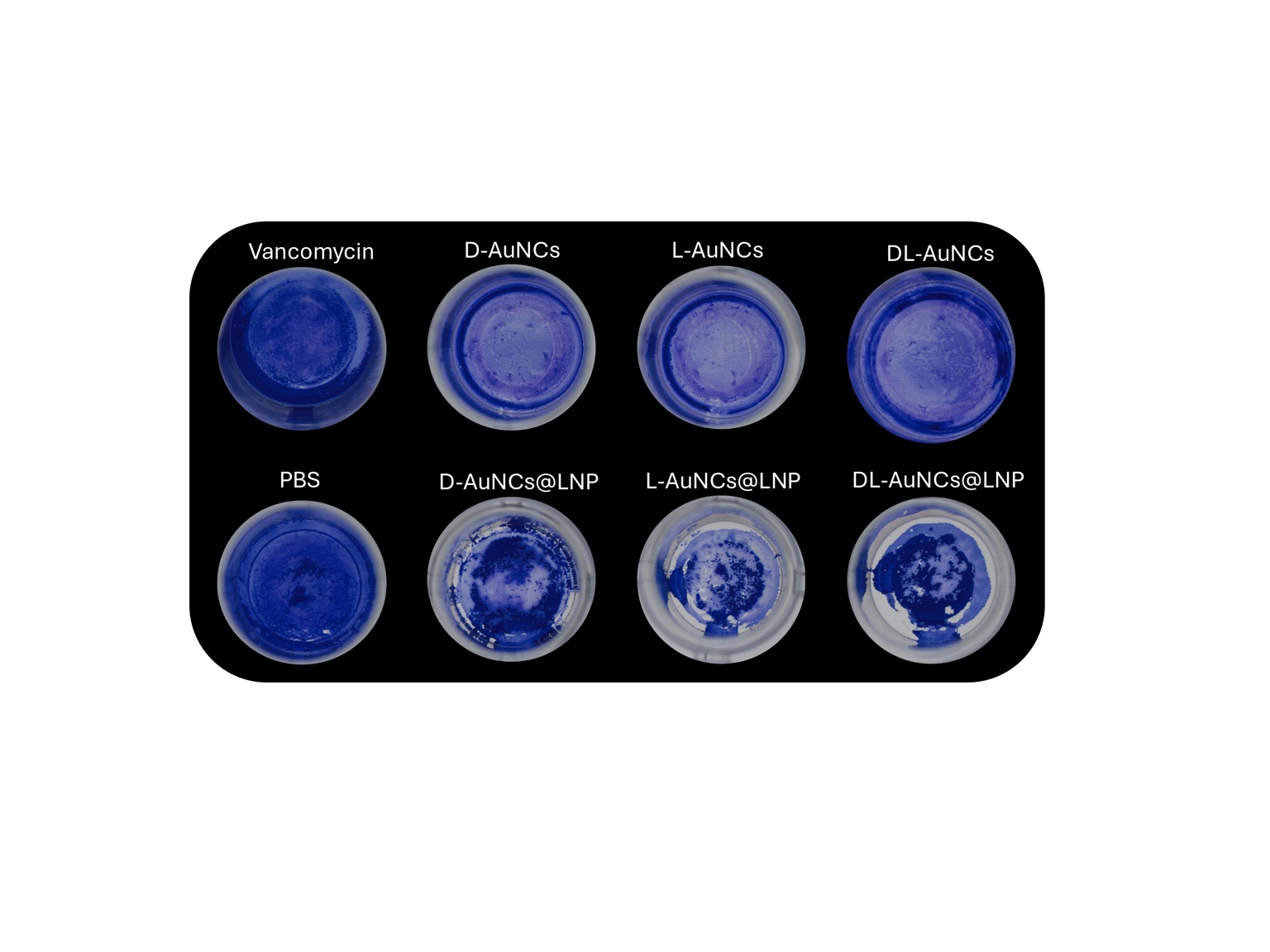

**Figure S4. Representative crystal violet (CV) images of *S. aureus* USA300 biofilms after treatment with chiral AuNCs or AuNCs@LNP at 75 μg/mL Au.** 24 h biofilms were treated for 24 h with PBS, vancomycin, D-, L-, or DL-AuNCs, or the corresponding AuNCs@LNP formulations at an Au-equivalent concentration of 75 μg/mL. Shown are representative wells after the standard CV workflow (wash–stain–wash) and prior to dye solubilization/absorbance quantification.

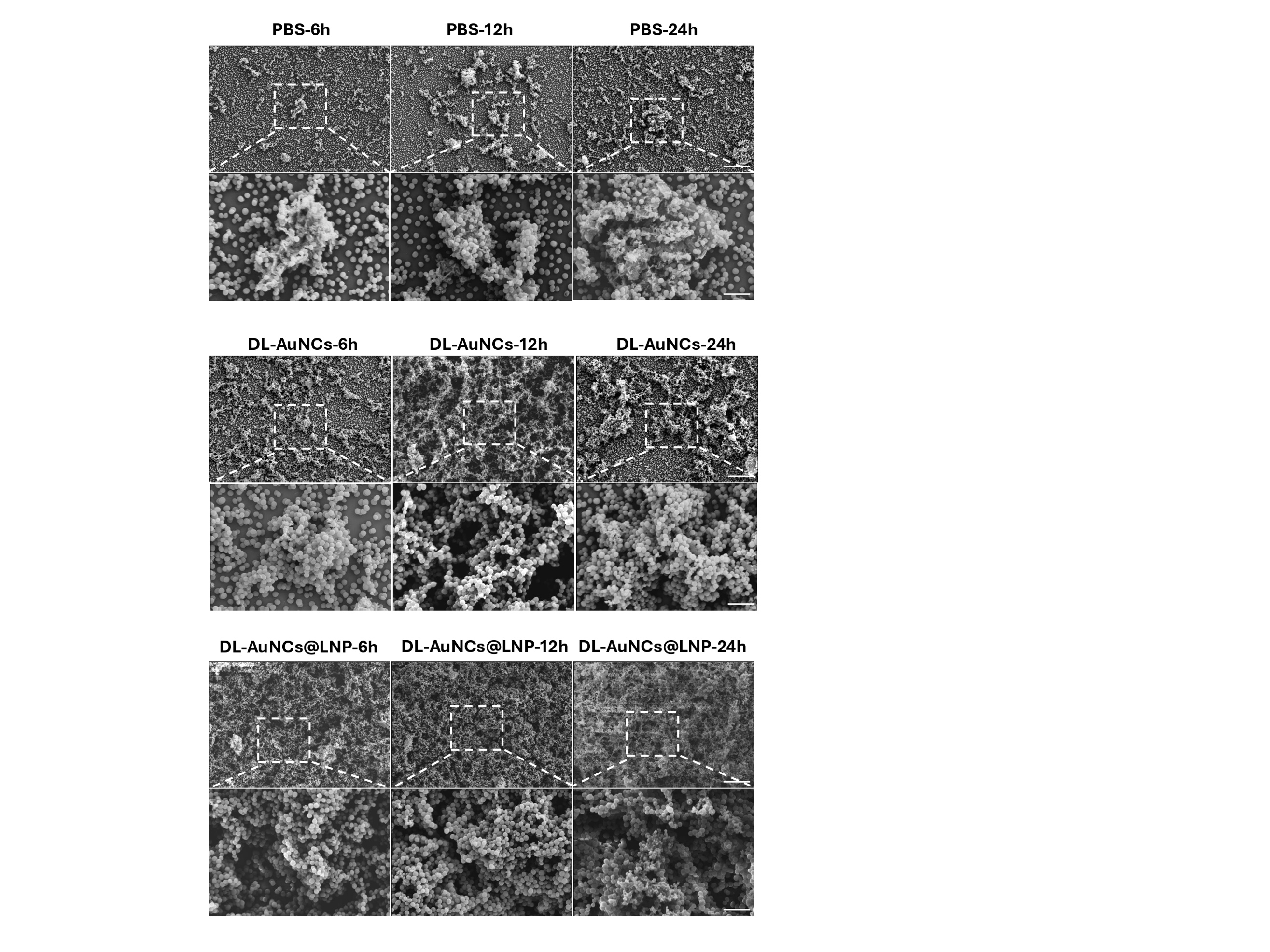

**Figure S5. Time-resolved SEM imaging highlights treatment-dependent remodeling of *S. aureus* USA300 biofilm architecture.** Representative scanning electron micrographs of established biofilms treated with PBS, DL-AuNCs, or DL-AuNCs@LNP (Au-equivalent 75 μg/mL) for 6, 12, or 24 h. PBS controls show structured surface coverage with intercellular connectivity consistent with an EPS-rich architecture. DL-AuNC treatment remodels the biofilm surface, whereas DL-AuNCs@LNP produces a distinct morphology with reduced obvious EPS-like bridging and abundant small spherical features interspersed among cells, consistent with pronounced architecture disruption in the formulated group. Upper panels show overviews (scale bars, 200 μm) and lower panels show higher-magnification views of the boxed regions (dashed outlines, scale bars, 5 μm).
